# Using gas exchange measurements to monitor growth, energy expenditure, and body composition of black soldier fly larvae (*Hermetia illucens*) on diets varying in protein-to-carbohydrate ratio

**DOI:** 10.1101/2025.09.08.674872

**Authors:** M.L. Schøn, K. Jensen, M. Schow-Madsen, M. Gold, A. Mathys, T.M. Schou, I.E. Berggreen, J.V. Nørgaard, J. Overgaard

**Affiliations:** Monogastric Nutrition, Department of Animal and Veterinary Sciences, Aarhus University, Blichers Allé 20, 8830 Tjele, Denmark; Sustainable Food Processing Laboratory, Department for Health Sciences and Technology, ETH Zurich, 8 Schmelzbergstrasse 9, 8092 Zürich, Switzerland; Zoophysiology, Department of Biology, Aarhus University, C.F. Møllers Allé 3, 8000 Aarhus C, Denmark; Enorm Biofactory A/S, Hedelundvej 15, 8762 Flemming, Denmark

**Keywords:** carbon dioxide, dietary macronutrient composition, energetic cost, growth monitoring, metabolism, oxygen, respiratory exchange ratio

## Abstract

Optimizing black soldier fly larvae (BSFL) production requires a better understanding of how diet composition shapes growth, nutrient allocation, and metabolic efficiency. This study investigated whether respiratory measurements could serve as predictive, non-invasive tools for BSFL performance on diets differing in protein-to-carbohydrate (P:C) ratio. Larvae were reared for seven days on four experimental diets, with growth, survival, gas exchange (O_2_ consumption, CO_2_ production, respiratory exchange ratio (RER)), and body composition (crude protein and lipid contents) measured across seven replicates per treatment. O_2_ consumption correlated strongly with final larval biomass, while RER was closely associated with protein and lipid deposition. Larvae on low-protein, high-carbohydrate diets accumulated proportionally more lipid, whereas protein-rich diets increased crude protein content but required higher energetic investment. This demonstrates that lipid-rich larvae grew more efficient in energetic terms per unit accumulated biomass than protein-rich larvae. Across diets, gas exchange variables reliably reflected growth dynamics, metabolic activity, and nutrient assimilation. Integrating respirometry into rearing systems could enable real-time monitoring of insect biomass yield and nutrient composition.

## Introduction

The world faces pressing environmental challenges that impact global food production, including climate change, soil degradation, habitat loss, and the depletion of natural resources (Foley *et al*., 2011). More sustainable food systems are essential to meet the increasing food and feed demand driven by population growth while simultaneously mitigating the impacts associated with greenhouse gas emissions, land-use, water scarcity, and food wastage (FAO, 2009; Foley *et al*., 2011). One component of this more sustainable food and feed production is the use of black soldier fly larvae (BSFL; *Hermetia illucens* (L.)) for converting low quality organic material into nutrient-rich biomass (Broeckx *et al*., 2021; Gold *et al*., 2020). This biomass can supplement traditional unsustainable protein sources (e.g. soybean and fishmeal) in livestock feeds (van der Heide *et al*., 2021) and pet foods (Bosch & Swanson, 2021), and as an added benefit, the remaining frass after harvest of larvae represents a high-quality fertilizer (Fuhrmann *et al*., 2022; Lomonaco *et al*., 2024).

The ability of BSFL to feed on a wide range of organic materials is a major reason why this species has become increasingly important commercially (Purkayastha & Sarkar, 2022). Nevertheless, BSFL growth rates, energy conversion efficiencies, and body compositions can vary considerably with diet quality and macronutrient balance (Barragan-Fonseca *et al*., 2021; Eggink *et al*., 2023), and with other conditions such as moisture content (Bekker *et al*., 2021) or temperature (Schøn *et al*., 2025). Diet composition, in particular the protein-to-carbohydrate (P:C) ratio and not just the total diet mass, has been shown to strongly affect larval growth and body composition (Cheon *et al*., 2022; Jensen *et al*., 2013; Lee & Jang, 2014; Rho & Lee, 2022). Studies have demonstrated that nutrient-deficient diets that deprive larvae of adequate protein or carbohydrate supply, as well as diets with unbalanced P:C ratios, reduce larval growth and alter nutrient deposition patterns (Barragan Fonseca *et al*., 2019; Berggreen *et al*., 2025; Eggink *et al*., 2023). As a result, differences in diet can lead to substantial variation in BSFL body composition, ranging from 32–58% crude protein and 15– 38% crude lipids on a dry matter (DM) basis (see review by Gold *et al*., 2018). Because variations in macronutrient composition (encompassing both the levels and specific forms of protein, fat, and carbohydrate) affect larval development, size, and composition, it is important to understand in more detail how these dietary factors influence metabolism and energy conversion during growth. Similarly, there is a growing interest in developing methods to monitor the progression of growth in actively rearing larvae, as repeated manual sampling and weighing is impractical, time consuming, and can disturb larval behaviour and substrate structure, potentially affecting growth dynamics.

Measurements of gas exchange (O_2_ consumption and CO_2_ production) as well as the respiratory exchange ratio (RER, VCO_2_/VO_2_) are powerful indicators of metabolism, energy expenditure, and nutrient assimilation under different dietary conditions (Parodi et al., 2020; Schøn *et al*., 2025; Talal *et al*., 2021). BSFL utilizes aerobic energy both for general maintenance of basal physiological processes, but also in association with the direct cost of growth and nutrient assimilation (Ferrannini, 1988; Talal *et al*., 2021) and studies have previously found a close association between O_2_ consumption or CO_2_ production and total larval mass gain (Eriksen, 2024; Schøn *et al*., 2025). However, this association may be affected by the rearing conditions, including the dietary conditions, as the energetics of growth is expected to differ depending on the ratio of lipid and protein assimilation and because different nutritional quality may affect the relative magnitude of maintenance metabolism versus growth metabolism in ectotherms (Goodrich *et al*., 2024; Laganaro *et al*., 2021). This may be further complicated by the fact that in the BSFL rearing system the diet and digestive tract microbiome also contribute to overall gas exchange rates (Bekker *et al*., 2021; Fuhrmann *et al*., 2025; Parodi *et al*., 2020). Nevertheless, continuous monitoring of gas exchange rates holds the potential to provide important information on growth and body composition in production systems. In this context, RER may be particularly relevant to examine, as this metric reflects the combined input from a range of metabolic processes related to both anabolism and catabolism. RER reflects the catabolism of macronutrients where oxidation of lipid, protein and carbohydrate result in RER of 0.7, 0.83 or 1.0, respectively (Schmidt-Nielsen, 1997). In addition, RER will increase as a consequence of lipogenesis from carbohydrates or amino acids (Ferrannini, 1988; Talal *et al*., 2021), and RER measurements therefore hold potential to indirectly monitor which macronutrients are metabolized and assimilated.

The main objectives of this study were: (*i*) to assess how altering diet macronutrient composition affects the growth, metabolism, and body composition of BSFL, and (*ii*) to determine whether gas exchange measurements during rearing are predictive of BSFL production output in terms of growth rate, quantity, and quality. To address these objectives, larval growth, gas exchange (O_2_ and CO_2_) and body composition (crude protein and lipid contents) were measured on BSFL reared on diets with varying P:C ratios.

## Materials and methods

### Black soldier flies

BSFL of 5-6 days upon hatching (expected 4^th^ or 5^th^ instar; Gligorescu *et al*. (2019)) were supplied from a commercial BSFL facility (Enorm Biofactory A/S, Flemming, Denmark). After collection at the factory, the larvae were transported (1 h) to Aarhus University, Denmark, where all experiments were performed. For each experimental replicate, 600 counted larvae with average individual mass of 5.0 ± 0.9 mg were placed in 1 L respirometer chambers (15 cm diameter, 6 cm height) each containing approximately 340 g of fresh diet. This resulted in a density of ∼1.76 larvae per g wet diet or ∼0.133 g dry diet per larva.

### Experimental protocol

The study investigated how BSFL metabolism is affected by dietary macronutrient composition by measuring gas exchange, growth, and body composition of BSFL during seven days of rearing on four experimental diets. Experiments were performed over four consecutive weeks using a blocked design where one or two replicates of each dietary treatment was included each week, resulting in seven replicates of each dietary treatment. The experimental system allowed for hourly recordings of O_2_ consumption and CO_2_ production. Larval growth was estimated daily by sub-sampling and weighing larvae from each replicate. At termination of the experiment, larvae were separated from the frass using wet sieving, and subsequently quickly dried with paper towels. Larvae were then weighed and photographed to count the number of survivors using CountThings (Dynamic Ventures Inc., California, US). All experiments were performed in a climate cabinet at 27 °C (Termaks KB8182, Bergen, Norway) and constant darkness. Larvae sampled at the termination and start of the experiment were stored at -20 °C before analyses of DM, crude lipid, and crude protein contents.

### Experimental diets

The ingredients and nutritional compositions of the four experimental diets are shown in Table 1. All diets contained commercial chicken feed (KylleKræs 1, Danish Agro, Galten, Denmark). The reference (REF) diet consisted of 100% chicken feed and was used as a high-protein diet control to monitor if the system supported growth and metabolism to a similar degree as previous studies (e.g., Berggreen *et al*., 2025; Eggink *et al*., 2023; Nguyen *et al*., 2015). The three semi-artificial diets where mixed to represent different ratios of protein to non-fibre carbohydrate (NFC) (P:C) within an industrially relevant range, and with the aim of identifying nutritionally limiting conditions in protein-restricted diets. Specifically, the P:C 1:3, P:C 1:5, and P:C 1:9 diets contained 35% chicken feed to ensure supplementation of all required micronutrients, and the remaining 65% was mixed to create semi-artificial diets of desired macronutrient contents following Berggreen *et al*. (2025). These treatment diets contained different proportions of a protein mix (casein and peptone, 1:1) and a NFC mix (sucrose and dextrin, 1:1) to achieve desired P:C ratios. A lipid mix (rapeseed oil, sunflower oil, and lard, 1:1:1) and a mix of additives including vitamins (Vanderzant modification vitamin mixture), minerals (Wessons salt mixture), fatty acids (linoleic acid and oleic acid, 4:1), cholesterol, and nipagin (1:6:1:1:1), as well as cellulose and agar were added at similar concentrations in the three mixed diets (Table 1). Cellulose and agar were added as structural components, and nipagin was added as a preservative. All experimental diets were prepared by mixing 320 g of agar solution (agar dissolved in boiling water and then cooled to 40 °C) with 80 g of pre-mixed dry diet ingredients. The diets were then left to solidify during which water evaporated, leaving 341.3 ± 6.3 g wet diet (∼77% water) in the chambers at the onset of the experiment.

**Table 1:** Macronutrient contents of the four experimental diets. The reference diet (REF) is a commercial chicken feed used as a reference diet supporting high growth. The three other diets are formulated in house and designed to represent different protein-to-carbohydrate (NFC) (P:C) ratios.

| **Dietary treatment** | **REF** | **P:C 1:3** | **P:C 1:5** | **P:C 1:9** |
| --- | --- | --- | --- | --- |
| Protein-to-carbohydrate ratio (P:C) | 1 : 3.2 | 1 : 3.2 | 1 : 5.0 | 1 : 9.4 |
| **Ingredients**<br>(% of dry mass) |  |  |  |  |
| Chicken feed (CF) | 97.6 | 34.1 | 34.1 | 34.1 |
| Protein mix | 0 | 10.4 | 5.2 | 0 |
| Carbohydrate mix | 0 | 33.3 | 38.6 | 43.7 |
| Lipid mix | 0 | 2.7 | 2.7 | 2.7 |
| Additive mix | 0 | 3.9 | 3.9 | 3.9 |
| Cellulose | 0 | 13.1 | 13.1 | 13.1 |
| Agar | 2.4 | 2.4 | 2.4 | 2.4 |
| **Diet composition**<br>(% of dry mass) |  |  |  |  |
| Protein | 19.9 | 17.3 | 12.1 | 7.0 |
| Carbohydrate (NFC) | 63.3 | 55.5 | 60.7 | 65.8 |
| Fibre (including cellulose and agar) | 5.6 | 16.7 | 16.7 | 16.7 |
| Lipid | 5 | 4.5 | 4.5 | 4.5 |
| Ash | 6.2 | 6.1 | 6.1 | 6.1 |

The diet composition is based on the ingredient list and the mass used. The non-fibre carbohydrate (NFC) fraction from chicken feed is estimated by subtracting protein, fat, fibre, and ash from the total dry mass.

### Gas exchange measurements

The open-flow respirometry setup and protocol for gas exchange measurements are described with greater detail in Schøn *et al*. (2025). Briefly, eight custom-built, airtight respiration chambers (one empty control chamber and seven experimental chambers) were placed in the climate cabinet. Each chamber was provided with a controlled and constant flow of room air (∼600 mL min^-1^) that continuously provided O_2_ and removed water vapor and CO_2_. The inlet air flow for each of the sequential chambers were regulated and recorded using a Flowbar-8 Mass Flow Meter System (Sable Systems). The recorded inlet flow of moist air was later corrected for water vapour dilution using a factor of 0.9877, thereby expressing the inlet flow (V_2I_) as dry air at ambient conditions, assuming a room air temperature of 21 °C, 50% RH, and an atmospheric pressure of 760 mmHg.

To measure VCO_2_ and VCCO_2_, air was sequentially sub-sampled at a rate of ∼200 mL min^-1^ using a SS-4 Sub-Sampler Pump (Sable Systems). Sequential shifting between chambers was performed using an eight-channel RM-8 Respirometry Flow Multiplexer (Sable Systems). The sampled air passed through a calcium chloride column for removal of water vapor, and through syringe filters (Acrodisc syringe filter, 25 mm, 0.2 µm Supor membrane, Pall Corporation) to remove particles, before passing through an O_2_ analyser (FC-2 Differential Oxygen Analyser (Oxzilla), Sable Systems) and a CO_2_ analyser (LI-850 Gas Analyzer, LI-COR Biosciences), respectively. The multiplexer shifted between each chamber every 7.5 minutes, allowing the readings to stabilize. Only the mean of the last 30 seconds of stable recordings was used, providing an hourly measurement for each chamber over the entire 7-day period. Unstable recordings of inlet flowrate, O_2_, or CO_2_ as well as outliers, were removed and replaced by interpolation or extrapolation from preceding and/or subsequent high-quality measurements, following the data quality assessment protocol described in the Supplementary Materials.

The rates of oxygen consumption (VCO_2_) and carbon dioxide production (VCCO_2_) (mL min^-1^) were calculated using Haldane-corrected equations from the inlet flow rate (VC_I_) and differences between inlet and outlet fractional gas concentrations, accounting for unequal inlet and outlet air volumes when RER ≠ 1 (Withers, 2001). Inlet gas fractions were estimated from the empty control chamber (F_IO_ and F_ICO_) and outlet fractions were measured from the experimental chambers (F_EO_ and F_ECO_) within the same hourly cycle.

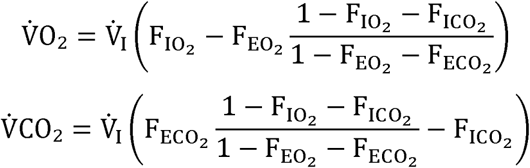

The total volumes of O_2_ consumed (VO_2_) and CO_2_ produced (VCO_2_) (L) for the 7-day experiments were calculated by summation of hourly measurements. The RER was calculated temporally as VCCOC/VCO_2_ and cumulatively as total VCO_2_/total VO_2_.

### Growth analysis

Respiration chambers with growing larvae were daily removed briefly from the temperature cabinet to track changes in total mass (frass and larvae) between days. On average the replicates lost 27.3 ± 3.4 g day^-1^, mainly due to removal of water vapor and CO_2_. During this procedure, 15 randomly selected larvae were quickly collected and weighed to assess average larval mass within each replicate. Three of the 15 larvae were sampled and frozen, and the other 12 larvae were returned. Thus, during the experiment 18 larvae were removed from each replicate. At the termination of the experiment all remaining larvae were counted and weighed to enable calculation of survival and total mass gain.

The total daily larval mass was estimated by multiplying the average daily larval mass with the estimated number of live larvae on each day. The estimate of live larvae accounted for the daily removal of 3 larvae and used an assumption that mortality was evenly distributed across the 7-day rearing period. Specific growth rates (SGR) were calculated as the difference in the natural logarithm of average body mass between two time points divided by the elapsed time:

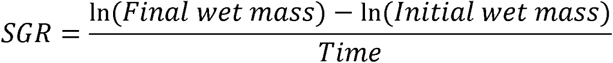

### Water, lipid, and protein content

At the termination of the experiment, all harvested larvae from each replicate were oven-dried at 60 °C for three to four days until they had reached a stable mass. Water content was calculated from the difference between wet mass and DM, and sub-samples of dried larvae were taken to estimate lipid and protein contents. To obtain crude lipid content, samples of 1.76 ± 0.01 g dried larvae were weighed before they were washed 5 times in 25 mL petroleum ether to remove the lipid fraction (Jensen *et al*., 2010). Each round of washing lasted 48 h and after the final round the defatted samples were again oven-dried at 60°C until stable mass to allow for estimation of the crude lipid fraction (calculated as the difference between initial DM and lean DM). Nitrogen analysis of defatted, finely ground sub-samples (4.2–7.2 mg per sample) was performed using a combustion analyser (Na 2000, Italy). The proportional crude protein content was then estimated by multiplying the measured nitrogen content by a factor of 4.76, a standardized conversion factor for the nitrogen-to-protein ratio in BSFL (Janssen *et al*., 2017). Total protein was estimated from lean DM and expressed as a fraction of the total DM (before lipid removal).

### Analysis of gross cost of growth and assimilation

To examine the relationship between energy consumption and energy assimilation, mass gain was converted into energy equivalents based on mass gain of lipid and protein. For these calculations an assumption of 37 kJ per g lipid and 17 kJ per g protein as energy conversion factors was used (FAO, 2002), respectively. Further, to assess the energy used for aerobic metabolism VO_2_ was converted to energy expenditure using a standard caloric conversion factor of 20.5 kJ per L O_2_ (Schmidt-Nielsen, 1997).

### Data analysis

Linear mixed-effects models (LMMs) were used to evaluate the effects of diet on parameters related to larval growth, gas exchange, and larval body composition. The four diet treatments were included as fixed effects, and experimental week was included as a random factor (model structure: Diet + (1 | Week)) to account for the observed small week-to-week baseline variations. Model assumptions were assessed by visual inspection of residual Q–Q plots and by residual diagnostics, with normality and homogeneity of variances evaluated using Shapiro–Wilk and Levene’s tests, respectively. Given the limited sample size per diet and week, these tests were interpreted cautiously and used as supplementary diagnostics rather than strict criteria. For a few parameters, minor deviations in residual diagnostics were observed. However, considering the low sample size, no substantial violations of model assumptions were detected, and the mixed-effects models were considered appropriate for interpretation of the data. Post-hoc comparisons among diet treatments were conducted using estimated marginal means with Sidak adjustment for multiple testing. Statistical significance was set at p < 0.05. Linear regressions were used to evaluate the relationships between final larval DM and VO_2_, as well as between assimilated energy from lipid and protein and the estimated aerobic energy expenditure. Pearson correlation coefficients were calculated to assess the strength and direction of these relationships. In selected analyses, linear models were constrained to pass through the origin (0,0) to test the assumption of a direct proportional relationship between the dependent and independent variables. Data processing and visualization were performed in RStudio version 2024.04.2 with R version 4.3.2 (R Core Team, 2023; RStudio Team, 2023).

## Results

Larval growth, gas exchange, and body composition varied by diet over the experimental period. Because larvae were supplied weekly, the experiments followed an unavoidable block structure, with each of the four experimental weeks representing a distinct batch. For some parameters, modest variation among weeks was visible in the data (see example in Figure S4). To account for this structure, experimental week was included as a random factor in the statistical models (LMMs), allowing diet effects to be estimated while accounting for week-to-week baseline variation. Diet consistently explained a large share of the variation across all response variables (Table S1). Accordingly, for all response variables except survival rate, diets differing in P:C ratio exerted a significant effect after accounting for week-to-week variation, and this biologically relevant effect is therefore the primary focus of the results and discussion presented below.

### Effects of diet on growth and body composition of BSFL

Larvae survived well and developed successfully on all diets, with average survival rates ranging from 89–97% and with no significant difference among diets (Table 2). While all diets supported substantial larval growth, the final mass of larvae was significantly different across diets after the 7-day growth period (Figure 1A). Total larval yield was highest when reared on the REF diet, followed by the mixed diets with higher protein fraction (Figure 1A, Table 2). The difference in larval growth between diets was also evident from our analysis of specific growth rates (SGR) (Figure 1B, Table 2). From the SGR (Figure 1B), it is possible to see the change in estimated total biomass between two consecutive days, where the biggest increase for P:C 1:3, P:C 1:5, and P:C 1:9 diets was found to be from day 3 to 4, and thereafter a more moderate increase (and even slight decrease for the REF diet) in SGR per day.

**Figure 1:**
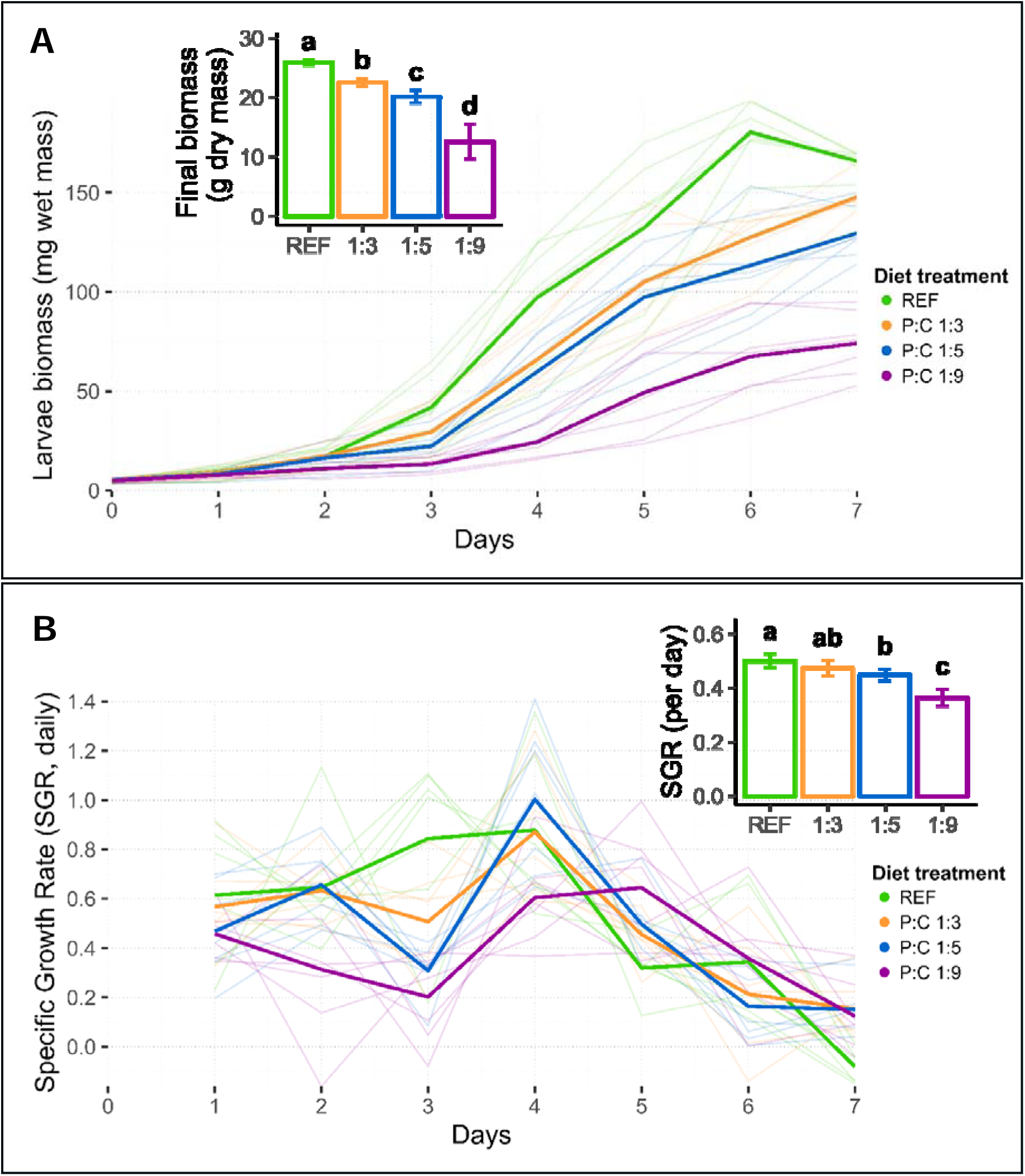
Effect of diets with different protein-to-carbohydrate ratios on total growth and estimated daily specific growth rate (SGR). A) Average individual larvae mass (mg wet mass) over time, where the insert shows the final total dry larvae biomass at the termination of the experiment (approximately 600 larvae). B) Daily estimates of SGR, calculated from the increase in average body mass estimated between two consecutive days, where the insert shows SGR calculated for the entire growth period. Bold lines represent treatment means, while semi-transparent lines show the individual replicates (n=7). Inserted bar plots show treatment means ± standard deviations, and dissimilar letters indicate significant differences based on linear mixed-effects models (p < 0.05, n = 7, Table S1).

**Table 2:** Survival, growth, body composition and gas exchange of black soldier fly larvae after 7 days of rearing on diets with different protein and carbohydrate contents.

|  | 5–6 day<br>old larvae* | REF | P:C 1:3 | P:C 1:5 | P:C 1:9 |
| --- | --- | --- | --- | --- | --- |
| **Growth performance** | At onset | After 7 days | After 7 days | After 7 days | After 7 days |
| Survival rate (%) |  | 97.0 ± 4.4%<sup>a</sup> | 94.9 ± 3.6%<sup>a</sup> | 92.9 ± 5.5%<sup>a</sup> | 88.8 ± 8.1%<sup>a</sup> |
| Larvae end biomass (g DM per replicate) | 0.78 ± 0.15 g | 25.9 ± 0.5 g<sup>a</sup> | 22.6 ± 0.7 g<sup>b</sup> | 20.1 ± 1.1 g<sup>c</sup> | 12.6 ± 3.0 g<sup>d</sup> |
| Specific growth rate (per day) |  | 0.50 ± 0.03<sup>a</sup> | 0.47 ± 0.03<sup>ab</sup> | 0.45 ± 0.02<sup>b</sup> | 0.36 ± 0.03<sup>c</sup> |
| Larvae percent dry matter (% DM) | 26.3% | 27.7 ± 1.5%<sup>a</sup> | 27.7 ± 0.8%<sup>a</sup> | 29.0 ± 2.2%<sup>a</sup> | 32.7 ± 1.5%<sup>b</sup> |
| Larvae crude protein (% of DM) | 40.9% | 31.5 ± 0.8%<sup>a</sup> | 32.5 ± 0.6%<sup>a</sup> | 25.5 ± 2.3%<sup>b</sup> | 19.8 ± 0.8%<sup>c</sup> |
| Larvae crude lipid (% of DM) | 10.0% | 24.3 ± 1.9%<sup>a</sup> | 27.5 ± 3.5%<sup>a</sup> | 39.2 ± 6.7%<sup>b</sup> | 52.7 ± 2.7%<sup>c</sup> |
| **Gas exchange measurements** |  |  |  |  |  |
| VCO<sub>2</sub> (L, total for 7 days) |  | 19.6 ± 1.2 L<sup>a</sup> | 16.9 ± 1.1 L<sup>b</sup> | 16.2 ± 1.5 L<sup>b</sup> | 10.8 ± 2.2 L<sup>c</sup> |
| VO<sub>2</sub> (L, total for 7 days) |  | 17.3 ± 1.3 L<sup>a</sup> | 13.4 ± 1.3 L<sup>b</sup> | 12.0 ± 1.4 L<sup>c</sup> | 7.3 ± 1.6 L<sup>d</sup> |
| RER |  | 1.13 ± 0.02<sup>a</sup> | 1.26 ± 0.05<sup>b</sup> | 1.35 ± 0.05<sup>c</sup> | 1.49 ± 0.09<sup>d</sup> |

\* Samples of 5-6-day old seed larvae were sampled for analysis at the onset of the experiment. P:C = Protein-to-carbohydrate ratio; DM = dry matter; V = volume; RER = respiratory exchange ratio, calculated from total VCO<sub>2</sub>/VO<sub>2</sub> over the 7-day experiment. All values are mean ± standard deviation and dissimilar letters indicate significant differences based on linear mixed-effects models (p < 0.05, n = 7, Table S1).

The larvae reared on different diets had significantly different body compositions at harvest (Table 2). Larvae reared on the P:C 1:9 diet had significantly lower water content compared to those from the other three diets. While the proportions of crude protein and crude lipid in the DM did not differ significantly between REF and P:C 1:3, they were significantly higher than found in larvae reared on diets with lower protein content (P:C 1:5 and P:C 1:9) (Table 2). As the protein content increased in the experimental diets, larval crude protein levels rose and lipid levels dropped, resulting in larvae from the low protein, high carbohydrate diets exhibiting significantly elevated proportional lipid fractions and reduced protein fractions (Table 2).

### Effects of diet on gas exchange and metabolism

The dynamic changes in VCOC and VCCO_2_ followed a broadly similar pattern (Figure 2A and Figure S1), both showing that gas exchange rates generally increased over the 7-day growth period but with different temporal progressions and distinct patterns depending on dietary treatment. For the commercial high-protein diet, REF, the VCO_2_ increased slowly during the first two days, after which the rate accelerated. VCO_2_ spiked at three distinct time points: between day 1-2 (0.6 mL O_2_ min^-1^), day 3-4 (2.2 mL O_2_ min^-1^), and day 5-6 (4.6 mL O_2_ min^-1^), before VCO_2_ declined considerably during the last 1.5 days of the experiment. A somewhat similar pattern was found for the P:C 1:3 and P:C 1:5 diets, but here the spikes occurred later and were more moderate. P:C 1:9 displayed an even more delayed and reduced response with a small spike around day 1.5-2.5, another by day 5.5, with indications of a third increase in VCO_2_ at the end of the experiment. Total VO_2_ differed significantly among all diet treatments, with the REF diet showing the highest O_2_ consumption, followed by progressively lower values in the mixed diets with decreasing dietary protein content (Figure 2A, Table 2).

**Figure 2:**
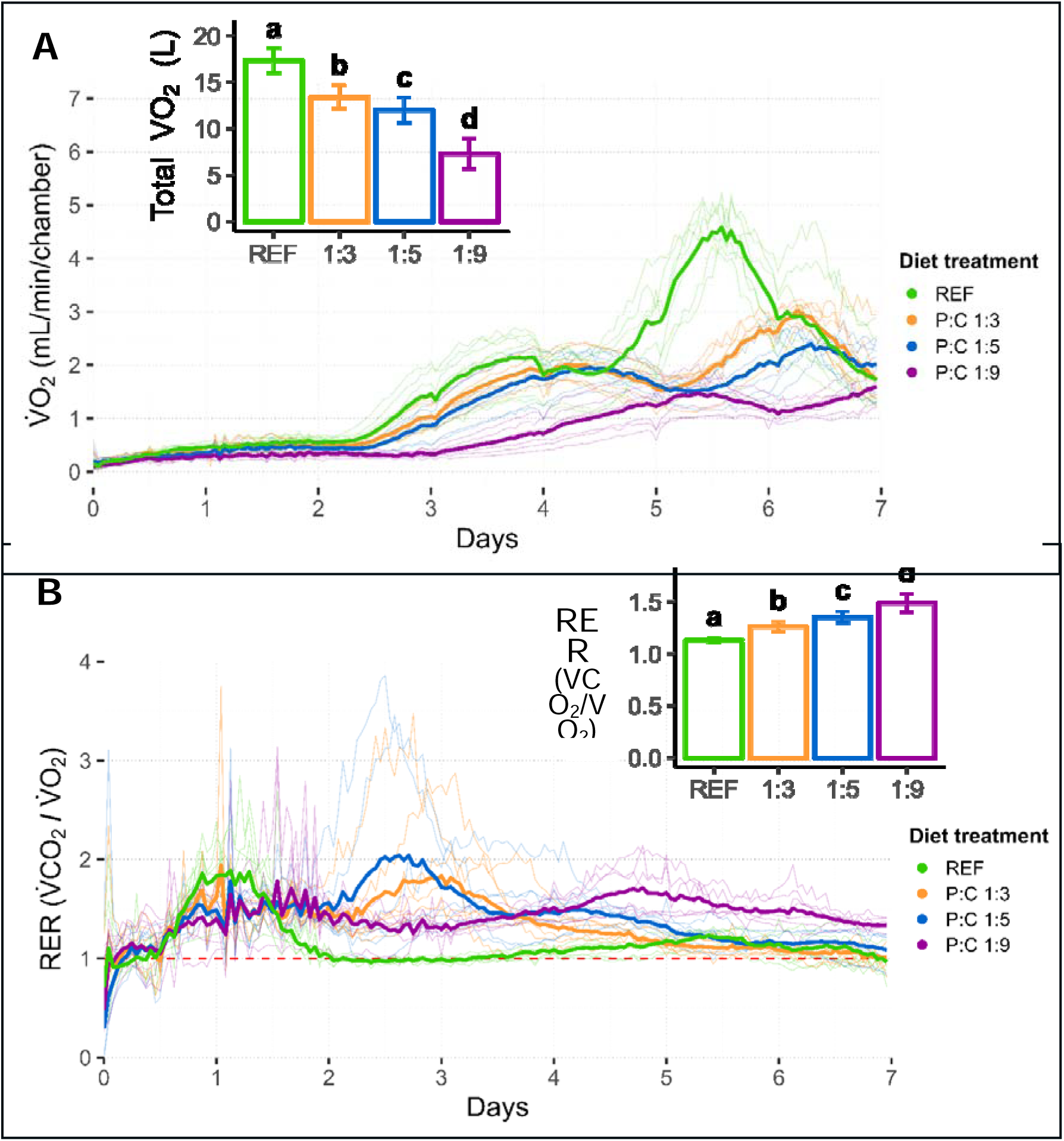
Effect of diet with different protein-to-carbohydrate ratios on gas exchange measurements. A) Change in O_2_ consumption rate (V̇O_2_, mL/min/chamber) over time, where the insert shows the total VO_2_ consumed for the entire period (approximately 600 larvae). B) Change in respiratory exchange ratio (RER) over time V̇CO_2_/V̇O_2_ (V̇ is the rate, mL/min), where insert show total measured RER VCO_2_/VO_2_(V_2_ is the volume, L) for each treatment. Bold lines represent treatment means, while semi-transparent lines show the individual replicates (n=7). Inserted bar plots show treatment means ± standard deviations, and dissimilar letters indicate significant differences based on linear mixed-effects models (p < 0.05, n = 7, Table S1).

RER values were >1 across diets when estimated from the total gas exchange over the 7-day growth period (Figure 2B, Table 2). RERs differ significantly among all four diets with lowest RER observed for the REF diet and increasing RER values with increasing dietary carbohydrate content.

### Respiratory variables as predictors of growth and body composition

Combining the data of growth and body composition with the data from gas exchange measures revealed several strong statistical correlations. There was a strong positive linear correlation between VO_2_ and final total larval DM when data was analysed across all diets (R² = 0.92, Figure 3A). This correlation persisted to be strong also when the linear regression model was forced through origo (0,0) (R² = 0.99, Figure 3A). As seen in Figure 3B, metabolic energy consumption per unit of assimilated energy as biomass (gross cost of growth) varied significantly among diets and resulted in a significant, albeit weaker, correlation (R² = 0.40, Figure 3B) suggesting that across diets 0.9 kJ was expended for each kJ assimilated. The cost of growth was significantly higher for the REF and lowest for the low protein diet (P:C 1:9). To examine this further, the correlation between overall energy consumption and protein energy assimilation was analysed. This analysis also found a strong positive correlation among all treatments rendering an overall energetic cost of 1.9 kJ per kJ of protein assimilated (R² = 0.87, Figure 3C). Importantly when analysed within each dietary treatment the cost of protein assimilation varied significantly between the semi-artificial diets (Figure 3C) such that protein energy assimilation was cheapest on the high protein diet (P:C 1:3) and more costly in the low protein diet (P:C 1:9).

**Figure 3:**
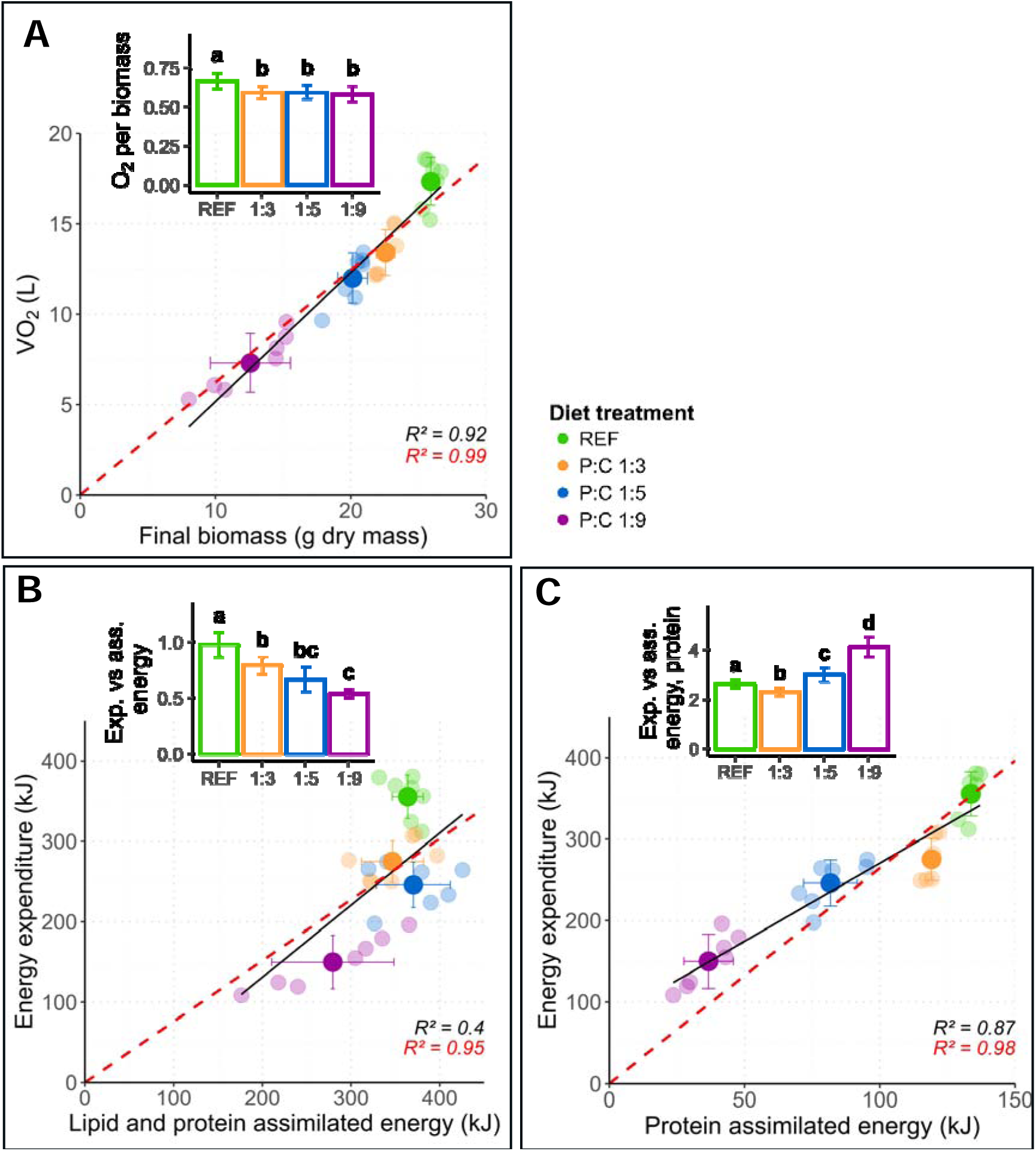
Analysis of the metabolic cost of growth and assimilation in larvae reared on diets with different protein-to-carbohydrate (P:C) ratio. A) Total oxygen consumption over 7 days as a function of final biomass across replicates reared on 4 different diets with varying P:C ratio. Correlation shows a linear relationship between the total volume of O_2_ consumed (L) and the final dry biomass (g). B) and C) show the energy expenditure of the system (using the energy equivalent of VO) in relation to the assimilated energy in the biomass (kJ) represented by gain in lipid and protein, or protein only. Bold points represent treatment means with error bars showing standard deviations, and semi-transparent points represent treatment replicates (n=7). Black lines represent a linear model for the replicates (n = 28), and red dashed lines represent the linear model when forced through origo (0,0) (regression statistics are reported in Table S2). Inserted bar plots show treatment means ± standard deviations, and dissimilar letters indicate significant differences based on linear mixed-effects models (p < 0.05, n = 7, Table S1).

Because RER reflects both catabolic and anabolic processes (i.e. lipogenesis) we hypothesized that this metric could reveal information about the body composition of growing larvae. Thus, lipid-rich larvae should be characterized by higher proportional CO_2_ contribution from lipogenesis than more lean larvae. As seen in Figure 4, RER was strongly correlated with both lipid and protein contents when analysed across larvae from all treatments (R² = 0.65 and R² = 0.64, respectively, Figure 4). When examining the data accounting for the experimental weeks, it becomes evident that week exerted a systematic influence on both RER and body composition, which helps explain the observed reversed within-diet correlations (Figure S4). This illustrates the relevance of including the experimental week as a random effect in the statistical analyses (Table S1).

**Figure 4:**
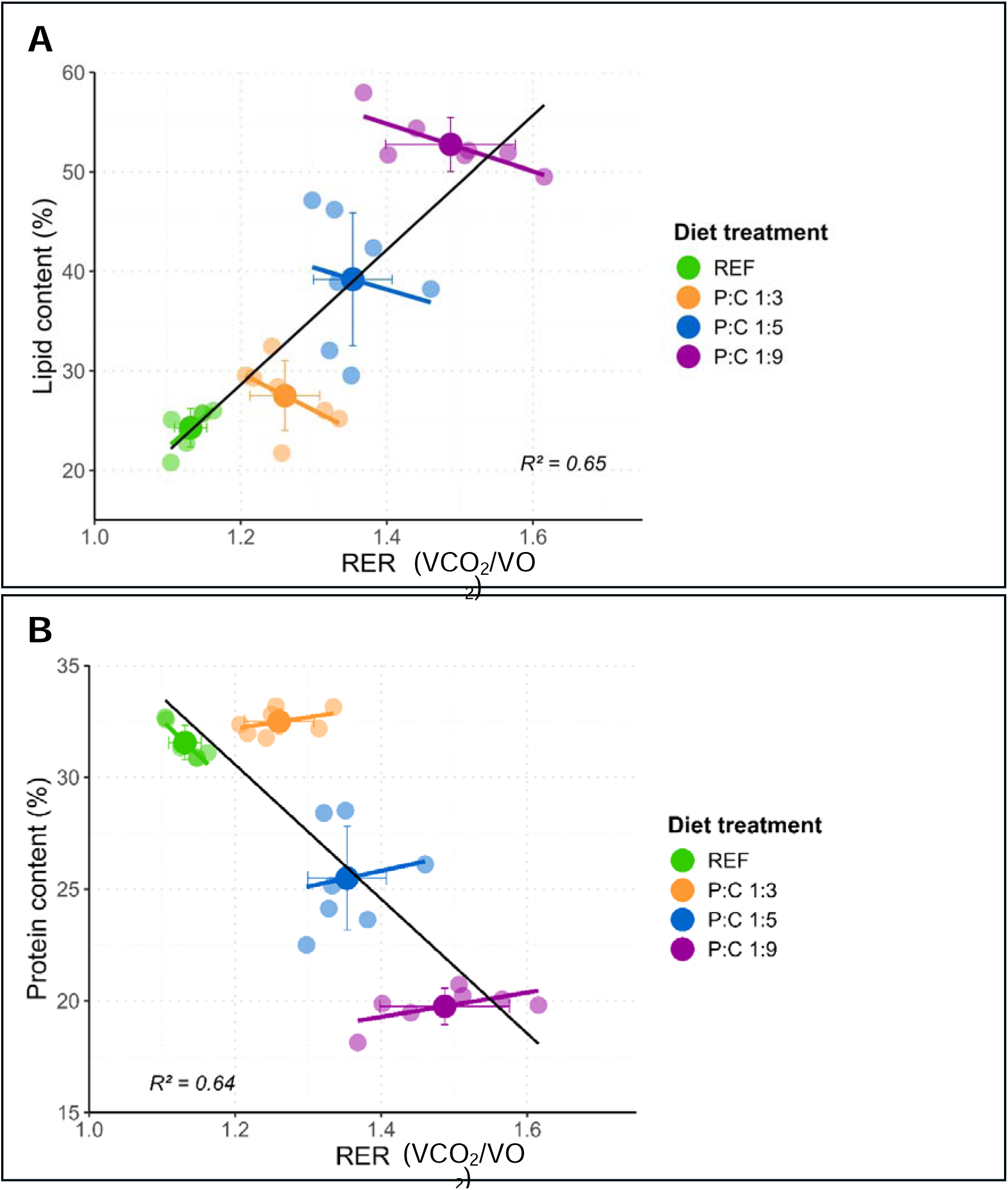
Effect of diets with different protein-to-carbohydrate ratios on larval body composition. Relationship between the measured respiratory exchange ratio (RER; VCO_2_/VO_2_) values and A) crude lipid and B) crude protein content (% of dry mass). Bold points represent treatment means with error bars showing standard deviations, and semi-transparent points represent treatment replicates (n = 7). The black lines represent a linear model for all the replicate data (n = 28). Linear models for each treatment group are shown in their respective colours (n = 7), and regression statistics are reported in Table S2.

## Discussion

The purpose of this study was to evaluate BSFL growth performance, metabolic activity, and energy conversion efficiency (energetic cost of growth) using respirometry as a monitoring tool. Four experimental diets with varying protein-to-carbohydrate (P:C) ratios were used; three semi-artificial diets with identical formulations except for varying P and C ratios, and a reference diet consisting of 100% chicken feed. Over a 7-day period we recorded larval growth, VCOC, VCCO_2_, and fractional content of crude protein and lipid of the final larvae. As expected, diets with a higher P:C ratio accelerated larval development, growth, and protein assimilation. Further, we found that the magnitude of oxygen consumption, CO_2_ production and RER provided useful and non-invasive measurements that correlated closely with larval output in terms of both developmental speed, total larval biomass and with regard to fractional lipid and protein content.

### Effects of dietary P:C ratio on growth and body composition of BSFL

The four experimental diets varied in P:C ratios but also in other aspects when the REF diet (chicken feed) was compared to the three semi-artificial diets. For example, we added a higher proportion of indigestible cellulose as a structural component to the semi-artificial diets, and the micronutrient concentrations also differed from that of the REF diet (see methods and Table 1). These differences may contribute to a slightly lower growth performance in the semi-artificial diets despite similar P:C ratio in the REF diet and the high-protein semi-artificial diet (P:C 1:3). Accordingly, the REF diet is primarily a valuable reference to earlier studies (Schøn *et al*., 2025; Schow-Madsen *et al*., 2026), whereas the three semi-artificial diets provide the best basis for direct comparison and assessment of the effects of P:C ratio.

Whether comparing all four diets or only the three semi-artificial diets, the present study confirms that high dietary protein fraction supports greater larval biomass and higher protein fraction in the larvae at harvest, whereas carbohydrate-rich diets increase the lipid fraction and produce smaller larvae (Table 1) (Barragan-Fonseca *et al*., 2021; Berggreen *et al*., 2025; Eggink *et al*., 2023; Gold *et al*., 2020). Protein-rich diet (P:C 1:3) resulted in 69.0% higher dry mass and 59.1% higher crude protein in the larvae compared to larvae from the carbohydrate-rich diet (P:C 1:9) (Table 1), and the P:C ratio of 1:3 has previously been termed “optimal” with regard to larval size and protein content (Eggink *et al*., 2023). In the P:C 1:9 diet the lipid fraction reached 52.7% of dry mass which is somewhat consistent with the >44% lipid fraction reported by Eggink *et al*., 2023 for similar P:C ratio. At the time of harvest on day 7 the larvae from the different dietary groups were likely at different developmental stages (see discussion below). This developmental difference may contribute to differences in body composition (Eggink & Dalsgaard, 2023) but harvesting larvae at different times introduce other limitations (Deruytter *et al*., 2025). Notably, Eggink *et al*. (2023) found that even when carbohydrate-rich diets (P:C 1:5 and P:C 1:9) were harvested to match developmental stage, they still produced larvae with significantly higher lipid content compared to larvae reared on protein-rich diets.

Some of the lipid assimilated in the larvae was likely converted directly from dietary lipid sources but our data support that *de novo* lipid synthesis from dietary carbohydrates sources was likely the main resource for larval lipid assimilation. Consistent with this view, larvae in high carbohydrate diets assimilated a much higher lipid fraction even though the dietary lipid fraction was almost identical in all diets (Table 1). Furthermore, the biochemical lipid compositions in BSFL larvae are known to deviate from that in the diet, suggesting that assimilated lipids are modified or generated *de novo* (Eggink *et al*., 2023; Giannetto *et al*., 2020). Finally, we recorded a large fractional surplus of CO_2_ (high RER) in larvae raised on carbohydrate-rich diets. As discussed below, this is consistent with a proportionally large CO_2_ production from carbohydrate-based lipogenesis (Schow-Madsen *et al*., 2026; Talal *et al*., 2021)

### Gas exchange rate as a correlate of growth and development in BSFL

In the present study we measured gas exchange (VCO_2_ and VCCO_2_) from a mixture of feed and larvae that likely included some contribution from diet-associated microbes. A recent study suggested that BSFL production on food waste involved substantial microbial CO_2_ production such that VCO_2_ increased fast and was maximised in the initial rearing phase while the total larval body mass was still small (e.g. Fuhrmann *et al*. 2025). However, the fibre-enriched food waste used in the study by Fuhrmann *et al*. (2025) likely included a much larger initial “microbial load” than the diets used in the present study that were freshly produced from dry ingredients and also included nipagen. Accordingly, we did not observe any large initial spike in gas exchange in our experiments. Instead the gas exchange intensity of all treatment groups tracked larval mass consistently with the asumption that gas exchange came mainly from larval metabolism. Furthermore, when RER was corrected to consider the CO_2_ presumed to originate from larval lipogenisis (see below), we arrived at RER estimates consistent with normal aerobic metabolism (Figure S3). Although we do not interpret the high RER measured across the entire 7-day growth period to be strongly biased from microbial CO_2_ production, we note that RER, on an hourly basis, was generally higher during the first 2-3 days (Figure 2B). This pattern suggests that microbial metabolism could play a proportionally larger role during the first days when the total metabolism of the system was low. Evenso, we consider that the total 7-day cumulative gas exchange was largely dominated by larval metabolism. Ideally, future experiments should be designed to carefully examine the partitioning between the magnitude of microbial and larval gas exchange and how it may change with time and diet type.

In the present study the larvae grew progressively on all diets (Figure 1A) and the temporal signal of metabolic intensity (Figure 2A and S1) showed increases and decreases that likely represent how the larval cohort in the respiration chambers progress through larval instars (Schøn *et al*., 2025; Schow-Madsen *et al*., 2026) (Figure 1A, 2A, and S1). Thus, moulting phases between instars temporarily reduces feeding and consequently lowers metabolic cost associated with growth (Gligorescu *et al*., 2019). The progression of “developmental peaks” differed among diets such that higher protein content accelerated development and shortened instar duration as indicated by closer peaks in metabolism (Figure 2A and S1). Although larval instar was not measured directly in the present study, these temporal profiles demonstrate the potential use of gas exchange to monitor developmental dynamics non-invasively in BSFL production systems (Schøn *et al*., 2025). For example, continuous measurements of gas exchange can identify when the cohort of larvae starts transitioning to the non-feeding 7th instar (large reduction in gas exchange rates) which will then indicate when larval biomass is approaching maximum. As an example, we observed for the REF diet that oxygen consumption rate peaked between day 5 and 6 (Figure 2A) followed by our recording of maximal mass on day 6 (Figure 1A). Larvae reared on P:C 1:3 and P:C 1:5 had their final metabolic peak (likely transition to 7th instar) between day 6 and 7, and for larvae reared on the P:C 1:9 diet the gas exchange rates were still increasing at day 7. These observations align well with Eggink *et al*. (2023) who found that carbohydrate-rich diet (P:C 1:9) prolonged larval development by approximately two days compared to diets with P:C ratios of 1:3.

### Energy expenditure and energy assimilation in BSFL

Aerobic energy consumption increases with growth in BSFL and other insects reflecting the cost associated with nutrient assimilation, tissue synthesis, and increased maintenance cost of the growing larval biomass (Eriksen, 2022, 2024; Ferral *et al*., 2020). While our experimental approach cannot directly partition the cost associated with either growth or maintenance (but see Eriksen, 2022, 2024) we confirmed that total oxygen consumption was a reliable and general predictor of final larval biomass across diets (Figure 3A). Comparing the four diets overall showed that O_2_ consumption per g biomass assimilated was slightly, but significantly, higher in the larvae on the REF diet (Figure 3A). This higher aerobic “cost” is likely explained by the observation that larvae on the REF diet reached maximal mass on day 6 while the cost for maintenance metabolism continued also on day 7 and because the REF diet supported growth of protein-rich larvae (see below).

In a more detailed analysis, we associated the total energetic expenditure (VO_2_ recalculated to kJ with a caloric coefficient of 20.5 J/ml) to either the total energetic gain in lipids and protein (Figure 3B) or to protein alone (Figure 3C). Larvae reared on the high-protein diet (P:C 1:3) expended ∼48% more metabolic energy per unit of lipid and protein than larvae reared on the high-carbohydrate diet (P:C 1:9) (Figure 3B). Thus, larvae with proportional high lipid allocation grew “cheaper” in energetic terms which is consistent with protein synthesis being inherently more energetically demanding than lipid accumulation (Goodrich *et al*., 2024; Laganaro *et al*., 2021; Talal *et al*., 2021). Conversely, we found the energetic cost per unit of protein gain was ∼44% lower in larvae reared on the P:C 1:3 diet compared to P:C 1:9 (Figure 3C). This pattern suggests reduced efficiency of protein deposition under protein-limited conditions while protein-rich diets seemingly facilitate a more energy-efficient protein assimilation. We caution, however, that isolating components of larval biomass assimilation and relating this to the overall energy use of the whole bioconversion system should be interpreted cautiously. This rough analysis does not directly partition growth- and maintenance-related costs (e.g. Eriksen, 2022, 2024) and protein-rich larvae likely may have higher maintenance costs per unit mass than lipid-rich larvae. Indeed, deviations between the unconstrained (black) and origin-forced (red) linear models in Figure 3C suggest that factors beyond a simple proportional relationship between metabolic cost and nutrient gain influence these patterns.

### Respiratory exchange ratio (RER) as a proxy of BSFL body composition

O_2_ consumption provides a good measure of total aerobic energy expenditure, while CO_2_ production arises from aerobic metabolism but potentially also from CO_2_ released upon biochemical conversion of dietary carbohydrates (or amino acids) into lipids (Laganaro *et al*., 2021; Talal *et al*., 2021). Specifically, RER (VCO_2_/VO_2_) from aerobic metabolism ranges from ∼0.7 for lipid oxidation to ∼1.0 for carbohydrate oxidation (Schmidt-Nielsen, 1997) but in fast-growing systems, lipogenesis can drive RER values above 1.0 due to CO_2_ production derived from lipid synthesis (Ferrannini, 1988; Talal *et al*., 2021; Parodi *et al*., 2020; Schøn *et al*., 2025). Furthermore, CO_2_ production may include contributions derived from microbial anaerobic metabolism which ads complexity to the analysis of CO_2_ production rates. Nevertheless, data from the present study support that RER can serve as a powerful and simple proxy for lipid assimilation and therefore also for the proportional lipid or protein fraction in the larvae at harvest (Figure 4).

Here we calculated cumulative RER based on total VO_2_ and total VCO_2_ over 7 days as this value captures the total gas exchange of the growth period and is less sensitive to experimental noise than hourly estimates. We found RER to be substantially higher for the high-carbohydrate diets that produced lipid-rich larvae (Figure 4, Table 2) and these differences in measured RER can almost perfectly be explained when considering CO_2_ production derived from lipogenesis (Figure 5). Thus, lipogenesis from carbohydrates generates 0.35 L CO_2_ per g of lipid (Ferrannini, 1988; Talal *et al.,* 2021) and we calculated a “corrected RER” assuming that 20% of BSFL lipid gain originated directly from dietary lipids while 80% was generated from carbohydrate-driven lipogenesis (see Figure S3). Across diets our estimate of “corrected RER” (ranging from 0.94-1.00) corresponds to aerobic catabolism based primarily on carbohydrate energy sources (Figure 5), but this calculation is obviously sensitive to our assumption regarding the proportion of carbohydrate-based lipogenesis. Notably, Talal *et al*. (2021) observed a similar pattern of high RER (>1.0) in locusts fed different P:C diets and the locust RERs were also “normalized” when corrected for lipogenesis (Figure S3). Although direct comparisons across insect species and dietary conditions should be made cautiously, we argue that RER can serve as a reliable proxy for BSFL fractional lipid or protein content. This was supported by data from the present study (Figure 4) as well as previous studies using comparable diets (Schøn *et al*., 2025; Schow-Madsen *et al*., 2026). However, it is clear that other sources of CO_2_ production (e.g. microbial metabolism) should be limited and controlled for this strong relationship to prevail.

**Figure 5:**
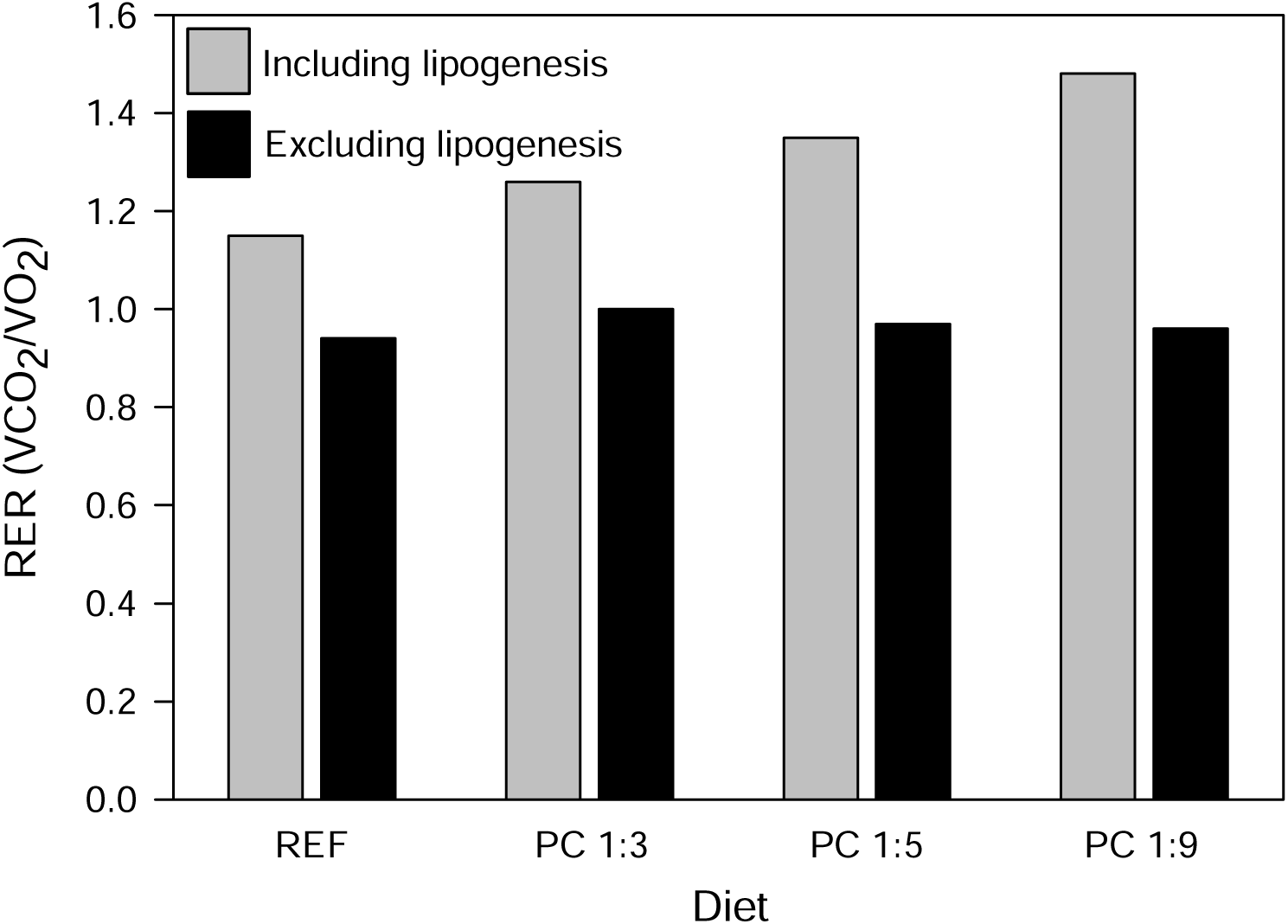
Effect of including or excluding the contribution of lipogenesis on the respiratory exchange ratio (RER). Bars show mean total measured RER (grey) and “corrected RER” (black), where CO_2_ produced from carbohydrate-derived lipogenesis is excluded. RER calculations were done using the assumption that 80% of the lipid mass gain comes from carbohydrate-driven lipogenesis and 20% comes from dietary lipid. Adjusting these assumptions would give slightly lower RER values at higher assumed carbohydrate-driven lipogenesis and slightly higher RER values at higher assumed contribution from dietary lipids. Comparing measured and “corrected” RER across diets shows that higher dietary carbohydrate content is associated with higher lipogenesis.

## Conclusion and perspectives

The present study focused on evaluating BSFL diet effects through the lens of gas exchange measurements and demonstrated how respiratory measurements can be applied as a predictive, non-invasive tool for monitoring growth performance, nutrient allocation, and energetic efficiency. Across diets with varying protein-to-carbohydrate (P:C) ratios, VCO_2_ varied consistent with developmental transitions, the total VO_2_ correlated strongly with final larval biomass, while RER reflected differences in protein and lipid deposition, confirming that dietary P:C ratio shapes both body composition and growth. Lipid-rich larvae on a high-carbohydrate diet grew more efficiently in energetic terms, but when evaluated in terms of protein assimilation this relationship was reversed highlighting practical implications for feed formulation and economic optimization. Thus, integrating O_2_ with CO_2_ monitoring in BSFL production facilities can link gas exchange to biomass yield and nutrient profiles, enabling real-time production monitoring to improve decision-making processes, resource use and ultimately valorization of organic side streams.

## Data availability

Data supporting this study will be made available upon publication.

## Declaration of competing interest

The authors declare that they have no known competing financial interests or personal relationships that could have influenced the work reported in this paper. This includes Moritz Gold who was employed at REPLOID Group AG in parallel to ETH Zurich during the writing phase of the presented research.

## Acknowledgements

This study was supported by the Danish Green Development and Demonstration Programme (GUDP), Projects EntoFeed 34009-21-1837 and FlyCloud 34009-20-1652.

## Supplementary material

**Figure S1:**
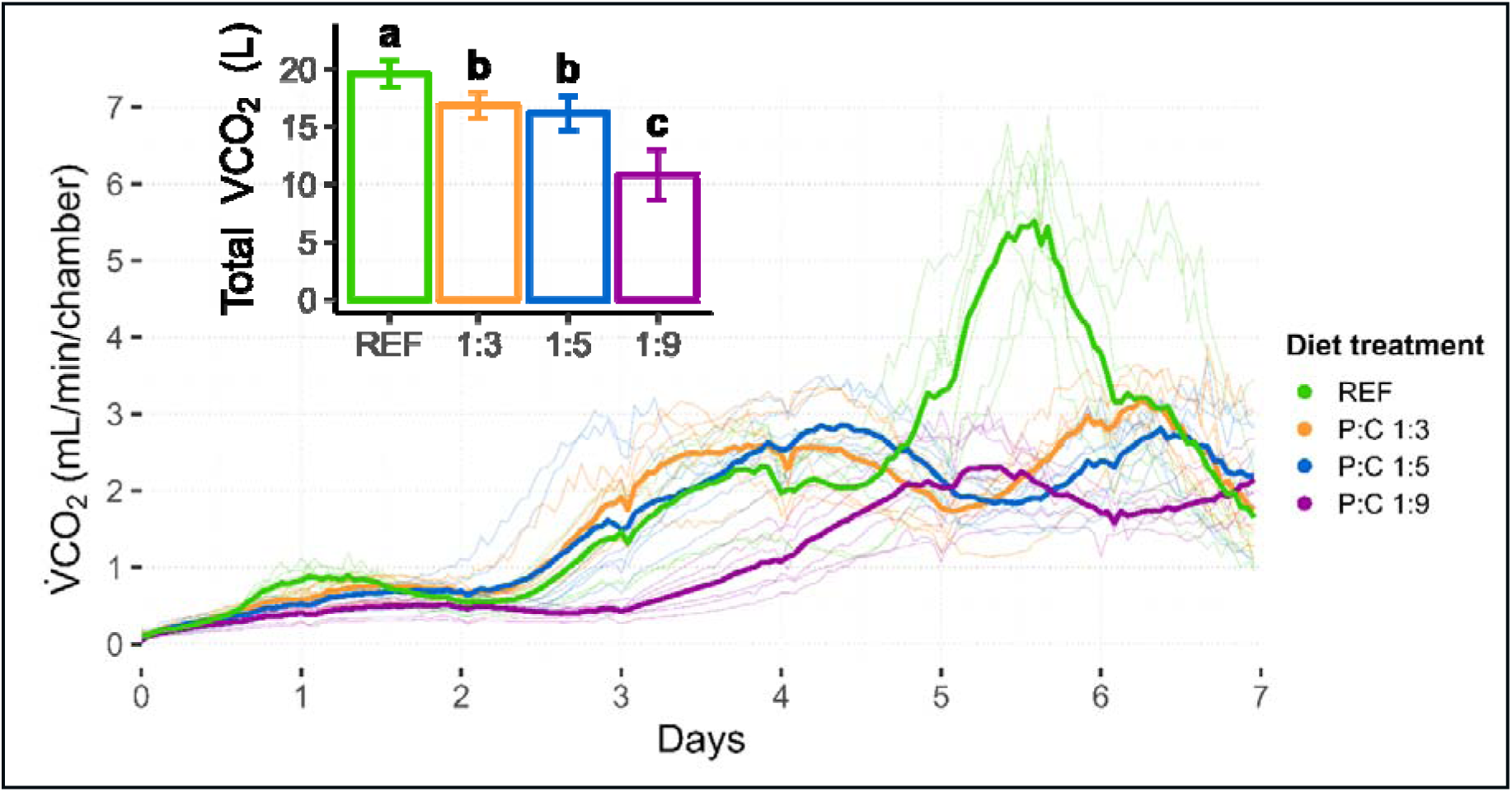
Effect of diet with different protein-to-carbohydrate ratios on CO_2_ production. Change in CO_2_ production rate (VCO_2_, mL/min/chamber) over time, where the insert shows the total VCO_2_ produced for the entire period (approximately 600 larvae). Bold lines represent treatment means, while semi-transparent lines show the individual replicates (n = 7). Inserted bar plots show treatment means ± standard deviations, and dissimilar letters indicate significant differences based on linear mixed-effects models (p < 0.05, n = 7, Table S1).

**Figure S2:**
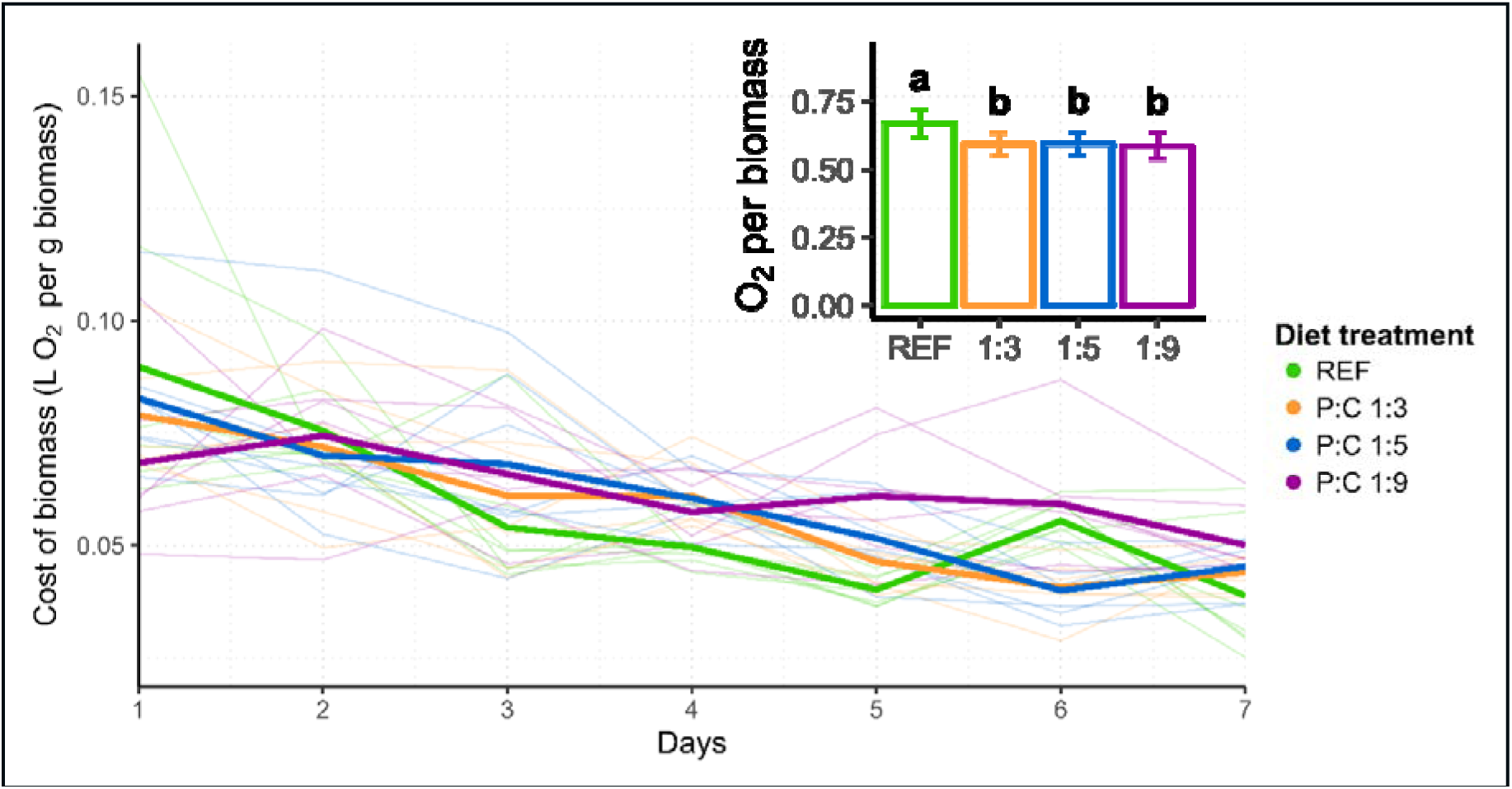
Effect of diet with different protein-to-carbohydrate ratios on the cost of biomass. Daily estimates of the metabolic costs of biomass, expressed as oxygen consumption (VO_2_, 24 h) per gram of total estimated larval biomass. Bold lines represent treatment means, while semi-transparent lines show the individual replicates (n = 7). Inserted bar plot show the total metabolic costs of final biomass with treatment means ± standard deviations, and dissimilar letters indicate significant differences based on linear mixed-effects models (p < 0.05, n = 7, Table S1).

**Figure S3:**
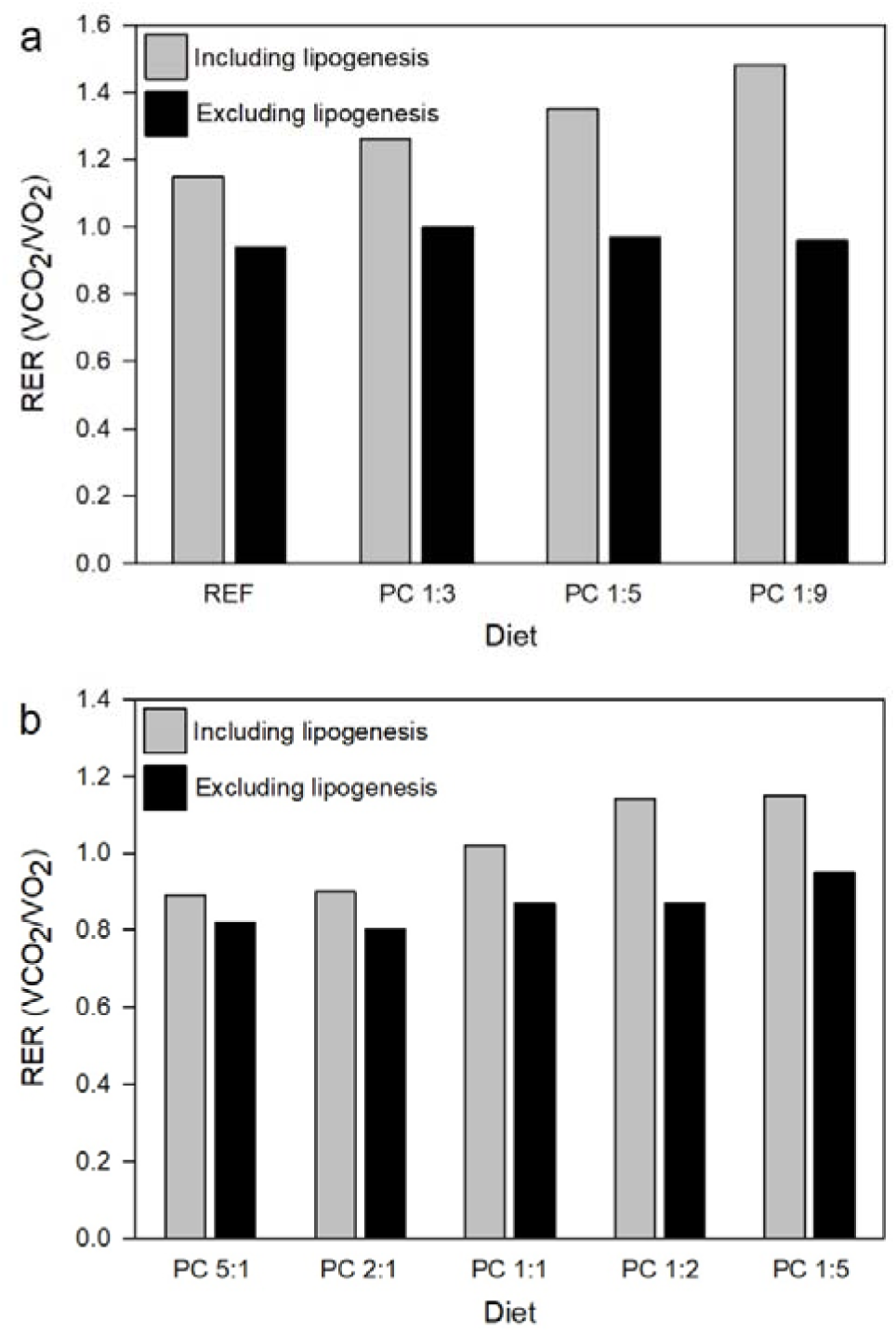
Effect of including or excluding the contribution of lipogenesis on the respiratory exchange ratio (RER) in a) the present study on black soldier fly larvae in wet diets and b) data adapted from the study by Talal *et al*. (2021) on grasshopper nymphs given dry diets. Bars show mean total measured RER (grey) and “corrected RER” (black), where CO_2_ produced from carbohydrate-derived lipogenesis is excluded. RER calculations for black soldier fly larvae were done using the assumption that 80% of the lipid mass gain comes from carbohydrate-driven lipogenesis and 20% comes from dietary lipid. Adjusting these assumptions would give slightly lower RER values at higher assumed carbohydrate-driven lipogenesis and slightly higher RER values at higher assumed contribution from dietary lipids. The result pattern resembles that of Talal *et al*. (2021) on dry diets where very limited microbial activity is expected, namely increasing proportions of lipogenesis at higher dietary carbohydrate levels, and fairly constant RER values close to 1 for black soldier fly larvae and 0.8 to 0.9 for grasshoppers when excluding the part of the respiratory output that is associated with lipogenesis. Comparing measured and “corrected” RER across diets and between the two experiments shows that higher dietary carbohydrate content is associated with higher lipogenesis, and that the same pattern is observed regardless of whether microbial activity was a likely contributor to overall metabolism or not. While some of the “excess” CO_2_ in wet diets may come from microbial fermentation, we propose that this analysis suggest that our measured RER mainly represent aerobic larval metabolism (including lipogenesis).

**Figure S4:**
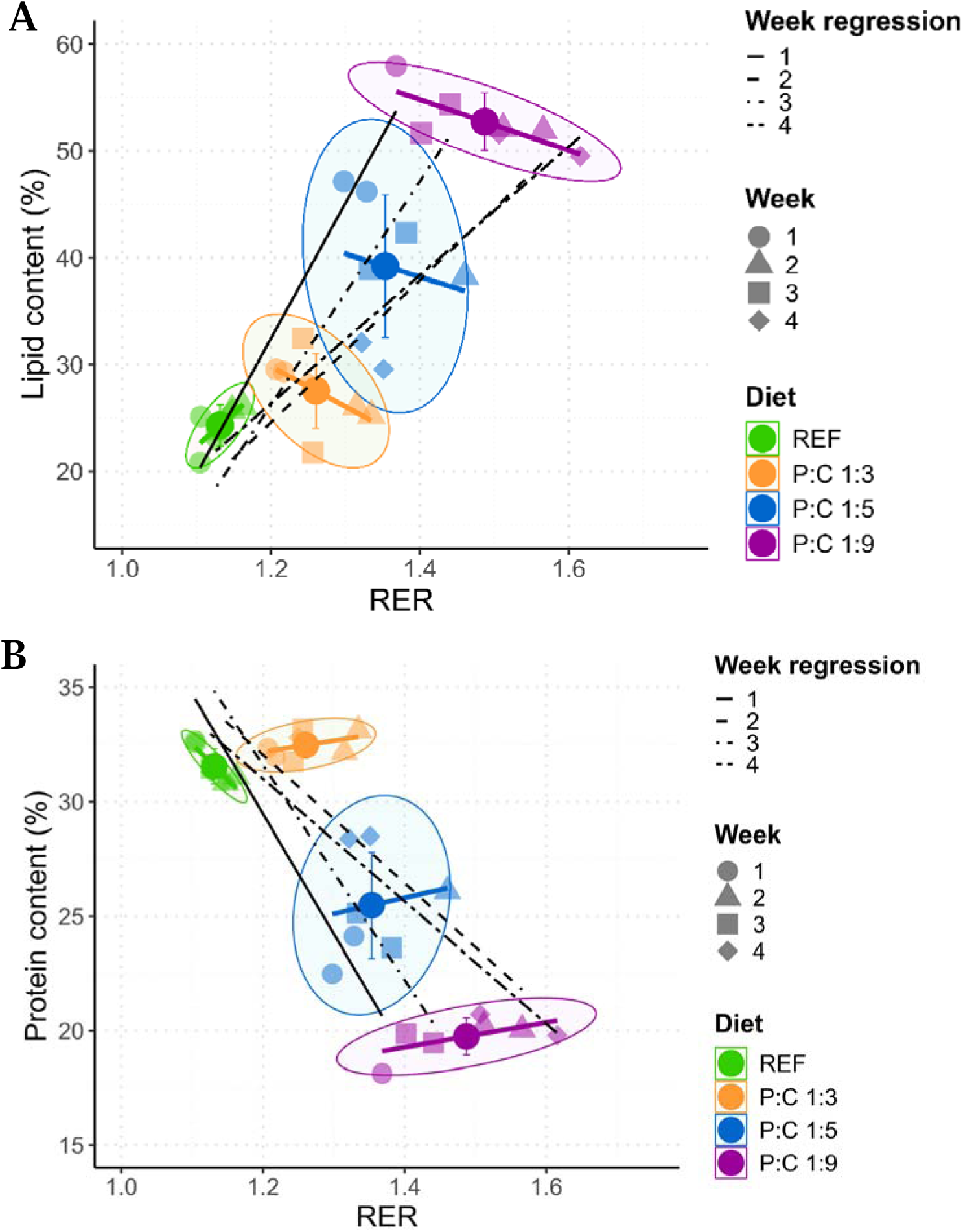
Week-specific variation in the relationship between RER and larval lipid and protein content. Relationship between the measured respiratory exchange ratio (RER; VCO_2_/VO_2_) values and A) crude lipid and B) crude protein content (% of dry mass). Bold points represent treatment means with error bars showing standard deviations, and semi-transparent points represent treatment replicates (n = 7). Symbols indicate the week in which each replicate was conducted. Black lines represent linear regressions fitted to replicate data within each experimental week (n = 7). Linear models for each treatment group are shown in their respective colours (n=7). Ellipses denote the 80% confidence regions for each diet. Regression statistics are reported in Table S2.

**Table S1:** Summary of linear mixed-effects model results for black soldier fly larvae growth, gas exchange, and body composition parameters. Diet was modelled as a fixed effect and experimental week as a random intercept (LMM: Diet + (1 | Week)). Pairwise post hoc comparisons were based on estimated marginal means with Sidak adjustment.

| Response variable | Diet<br>F-statistic | Diet<br>p-value | Week<br>variance | Post-hoc grouping |
| --- | --- | --- | --- | --- |
| Survival rate (%) | 2.73 | 0.07 | 1.93 | Not significant |
| Larval DM (g) | 126.83 | <0.001 | 1.08 | REF<sup>a</sup>, P:C 1:3<sup>b</sup>, P:C 1:5<sup>c</sup>, P:C 1:9<sup>d</sup> |
| Larval DM (%) | 30.97 | <0.001 | 1.38 | REF<sup>a</sup>, P:C 1:3<sup>a</sup>, P:C 1:5<sup>a</sup>, P:C 1:9<sup>b</sup> |
| Larval fat (% DM) | 89.08 | <0.001 | 4.31 | REF<sup>a</sup>, P:C 1:3<sup>a</sup>, P:C 1:5<sup>b</sup>, P:C 1:9<sup>c</sup> |
| Larval protein (% DM) | 161.13 | <0.001 | 0.25 | REF<sup>a</sup>, P:C 1:3<sup>a</sup>, P:C 1:5<sup>b</sup>, P:C 1:9<sup>c</sup> |
| Total VCO<sub>2</sub> (L) | 66.47 | <0.001 | 1.20 | REF<sup>a</sup>, P:C 1:3<sup>a</sup>, P:C 1:5<sup>b</sup>, P:C 1:9<sup>c</sup> |
| Total VO<sub>2</sub> (L) | 153.19 | <0.001 | 1.42 | REF<sup>a</sup>, P:C 1:3<sup>b</sup>, P:C 1:5<sup>c</sup>, P:C 1:9<sup>d</sup> |
| Specific growth rate (SGR) | 79.70 | <0.001 | 0.00 | REF<sup>a</sup>, P:C 1:3<sup>ab</sup>, P:C 1:5<sup>b</sup>, P:C 1:9<sup>c</sup> |
| Respiratory exchange ratio (RER) | 88.10 | <0.001 | 0.00 | REF<sup>a</sup>, P:C 1:3<sup>b</sup>, P:C 1:5<sup>c</sup>, P:C 1:9<sup>d</sup> |
| O<sub>2</sub> per biomass (L per g) | 6.25 | <0.001 | 0.00 | REF<sup>a</sup>, P:C 1:3<sup>b</sup>, P:C 1:5<sup>b</sup>, P:C 1:9<sup>b</sup> |
| CO<sub>2</sub> per biomass (L per g) | 8.57 | <0.001 | 0.00 | REF<sup>a</sup>, P:C 1:3<sup>a</sup>, P:C 1:5<sup>ab</sup>, P:C 1:9<sup>b</sup> |
| Energy exp./ass. (protein + lipid) | 31.65 | <0.001 | 0.00 | REF<sup>a</sup>, P:C 1:3<sup>b</sup>, P:C 1:5<sup>bc</sup>, P:C 1:9<sup>c</sup> |
| Energy exp./ass. (protein) | 98.28 | <0.001 | 0.04 | REF<sup>a</sup>, P:C 1:3<sup>b</sup>, P:C 1:5<sup>c</sup>, P:C 1:9<sup>d</sup> |

DM = dry matter; V = volume; RER is calculated from total VCO<sub>2</sub>/VO<sub>2</sub>; Diet treatments are REF, P:C 1:3, P:C 1:5, and P:C 1:9; P:C = Protein-to-carbohydrate ratio; Dissimilar letters indicate significant differences based on the linear mixed-effects models ( $p < 0.05$ , $n = 7$ ). Energy exp./ass. = the ratio of total metabolic energy expenditure to assimilated energy retained in larval biomass, based on protein and lipid gain (protein + lipid) or protein gain alone.

**Table S2:** Key statistics from the linear regression models shown in Figure 3, 4, and S4. Summary of the model formula, regression function, explained variance (R²), F-statistic, p-value, sample size (n), and Pearson correlation coefficient.

| Model illustration | Model formula | Factor | Regression function | $R^2$ | F-statistic | p-value | n | Pearson correlation coefficient |
| --- | --- | --- | --- | --- | --- | --- | --- | --- |
| Fig. 3A black | Total $VO_2$ (L) ~ Final larvae biomass (g DM) | | $f(x) = 0.7x + -2.0$ | 0.92 | 308.1 | < 0.001 | 28 | 0.96 |
| Fig. 3A red | Total $VO_2$ (L) ~ Final larvae biomass (g DM) + 0 | | $f(x) = 0.6x$ | 0.99 | 3362.6 | < 0.001 | 28 | 0.96 |
| Fig. 3B black | Energy expenditure (kJ) ~ Protein and lipid energy gain (kJ) | | $f(x) = 0.9x + -48.7$ | 0.40 | 17.2 | < 0.001 | 28 | 0.63 |
| Fig. 3B red | Energy expenditure (kJ) ~ Protein and lipid energy gain (kJ) + 0 | | $f(x) = 0.8x$ | 0.95 | 492.1 | < 0.001 | 28 | 0.63 |
| Fig. 3C black | Energy expenditure (kJ) ~ Protein energy gain (kJ) | | $f(x) = 1.9x + 78.5$ | 0.87 | 178.7 | < 0.001 | 28 | 0.93 |
| Fig. 3C red | Energy expenditure (kJ) ~ Protein energy gain (kJ) + 0 | | $f(x) = 2.6x$ | 0.98 | 1139.5 | < 0.001 | 28 | 0.93 |
| Fig. 4A black | Larvae lipid (% DM) ~ RER | | $f(x) = 67.5x + -52.4$ | 0.65 | 48.1 | < 0.001 | 28 | 0.81 |
| Fig. 4A and S4A diets | Larvae lipid (% DM) ~ RER | REF | $f(x) = 62.1x + -46.0$ | 0.51 | 5.2 | 0.07 | 7 | 0.71 |
| | | P:C 1:3 | $f(x) = -37.4x + 74.7$ | 0.26 | 1.8 | 0.24 | 7 | -0.51 |
| | | P:C 1:5 | $f(x) = -21.6x + 68.5$ | 0.03 | 0.2 | 0.71 | 7 | -0.17 |
| | | P:C 1:9 | $f(x) = -23.8x + 88.1$ | 0.61 | 7.9 | < 0.05 | 7 | -0.78 |
| Fig. 4B black | Larvae protein (% DM) ~ RER | | $f(x) = -30.1x + 66.6$ | 0.64 | 47 | < 0.001 | 28 | -0.80 |
| Fig. 4B and S4B diets | Larvae protein (% DM) ~ RER | REF | $f(x) = -31.2x + 66.8$ | 0.81 | 20.9 | < 0.01 | 7 | -0.90 |
| | | P:C 1:3 | $f(x) = 5.0x + 26.2$ | 0.17 | 1.1 | 0.35 | 7 | 0.42 |
| | | P:C 1:5 | $f(x) = 7.0x + 16.0$ | 0.03 | 0.1 | 0.73 | 7 | 0.16 |
| | | P:C 1:9 | $f(x) = 5.4x + 11.7$ | 0.35 | 2.7 | 0.16 | 7 | 0.59 |
| Fig. S4A weeks | Larvae lipid (% DM) ~ RER | Week 1 | $f(x) = 126.8x + -119.6$ | 0.92 | 56.0 | < 0.001 | 7 | 0.96 |
| | | Week 2 | $f(x) = 66.6x + -55.3$ | 0.77 | 16.9 | < 0.01 | 7 | 0.88 |
| | | Week 3 | $f(x) = 104.7x + -99.4$ | 0.82 | 22.4 | < 0.01 | 7 | 0.90 |
| | | Week 4 | $f(x) = 60.1x + -45.8$ | 0.88 | 36.7 | < 0.01 | 7 | 0.94 |
| Fig. S4B weeks | Larvae protein (% DM) ~ RER | Week 1 | $f(x) = -52.4x + 92.4$ | 0.82 | 22.2 | < 0.01 | 7 | -0.90 |
| | | Week 2 | $f(x) = -28.0x + 65.7$ | 0.68 | 10.5 | < 0.05 | 7 | -0.82 |
| | | Week 3 | $f(x) = -46.9x + 87.9$ | 0.79 | 19.2 | < 0.01 | 7 | -0.89 |
| | | Week 4 | $f(x) = -26.7x + 63.1$ | 0.86 | 31.3 | < 0.01 | 7 | -0.93 |

DM = dry matter; V = volume; RER = respiratory exchange ratio, calculated from total $VCO_2/VO_2$ over the 7-day experiment; Diets = REF, P:C 1:3, P:C 1:5, and P:C 1:9; Red lines represent the linear model when forced through origo (0,0).

## Protocol: Quality assessment of respirometry measurements

### Data collection

As described in the Methods, air from each of the eight respirometry chambers were sampled sequentially in repeated 7.5-minute cycles. When switching between each chamber, O_2_ and CO_2_ signals typically required a short washout period to reach steady state due to residual air in the system. Therefore, only the last 30 seconds of each interval were used to calculate a mean gas concentrations and flow rate.

### Manual screening

Measurements and data were visually inspected and excluded from the dataset if showing instability, drift, or unrepresentative fluctuations. See example in Figure S5 below.

### Low-signal exclusion

Measurements with gas exchange rates below 0.1 mL min ¹ after the first hour were excluded, as such low values was only observed when experiencing temporary loss of chamber airtightness.

### Automatic outlier detection

After manual and low-signal exclusions, an automatic outlier detection step performed in R Studio (RStudio Team, 2023) was applied to the remaining data. For each replicate’s time series data, the expected value at each time point was calculated as the mean of the three nearest valid measurements before and after this time point. A measurement was excluded as an outlier if it deviated by more than 20% from this mean. The first and last four measurements of each time series were not evaluated to avoid edge effects.

### Replacement of excluded measurements

Because O_2_ and CO_2_ concentration measurements are not independent, if one gas signal was considered unreliable, the paired gas value was also excluded. Measurements excluded by the data quality criteria described above were subsequently replaced by interpolation, or extrapolation where necessary, using the three nearest valid measurements before and/or after the missing data point.

A visual example of the quality assessment is shown in Figure S6.

**Figure S5:**
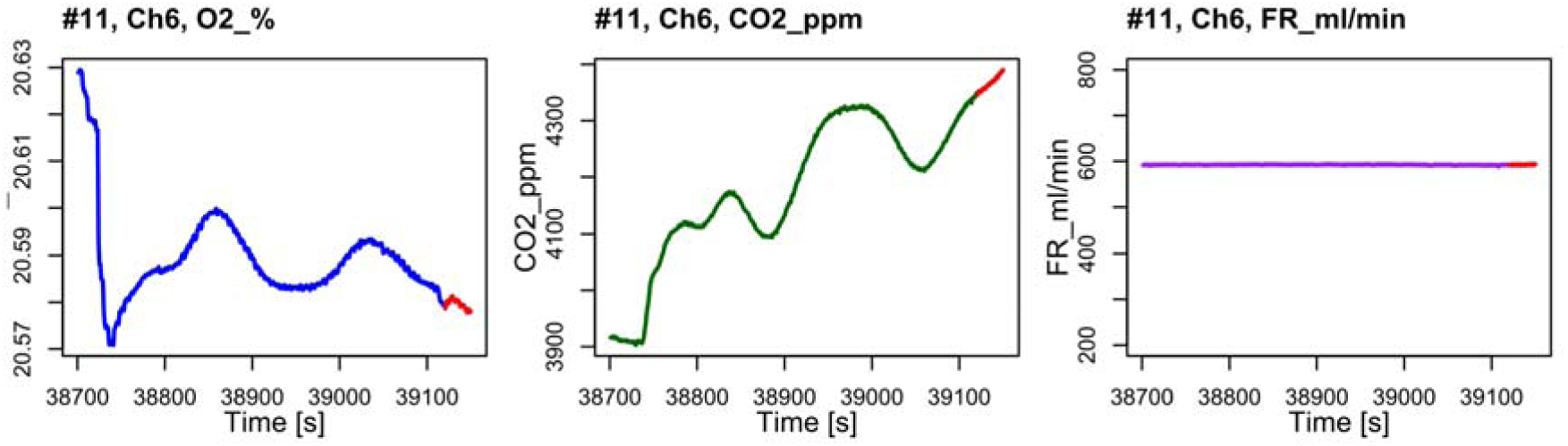
Example of an unstable measurement of gas concentrations (O_2_ (%) and CO_2_ (ppm)) during the final 30 seconds of a cycle. The graphs show the 7.5-minute recording for chamber 6 (P:C 1:3 diet) in experimental week 2, day 6, hour 11.

**Figure S6:**
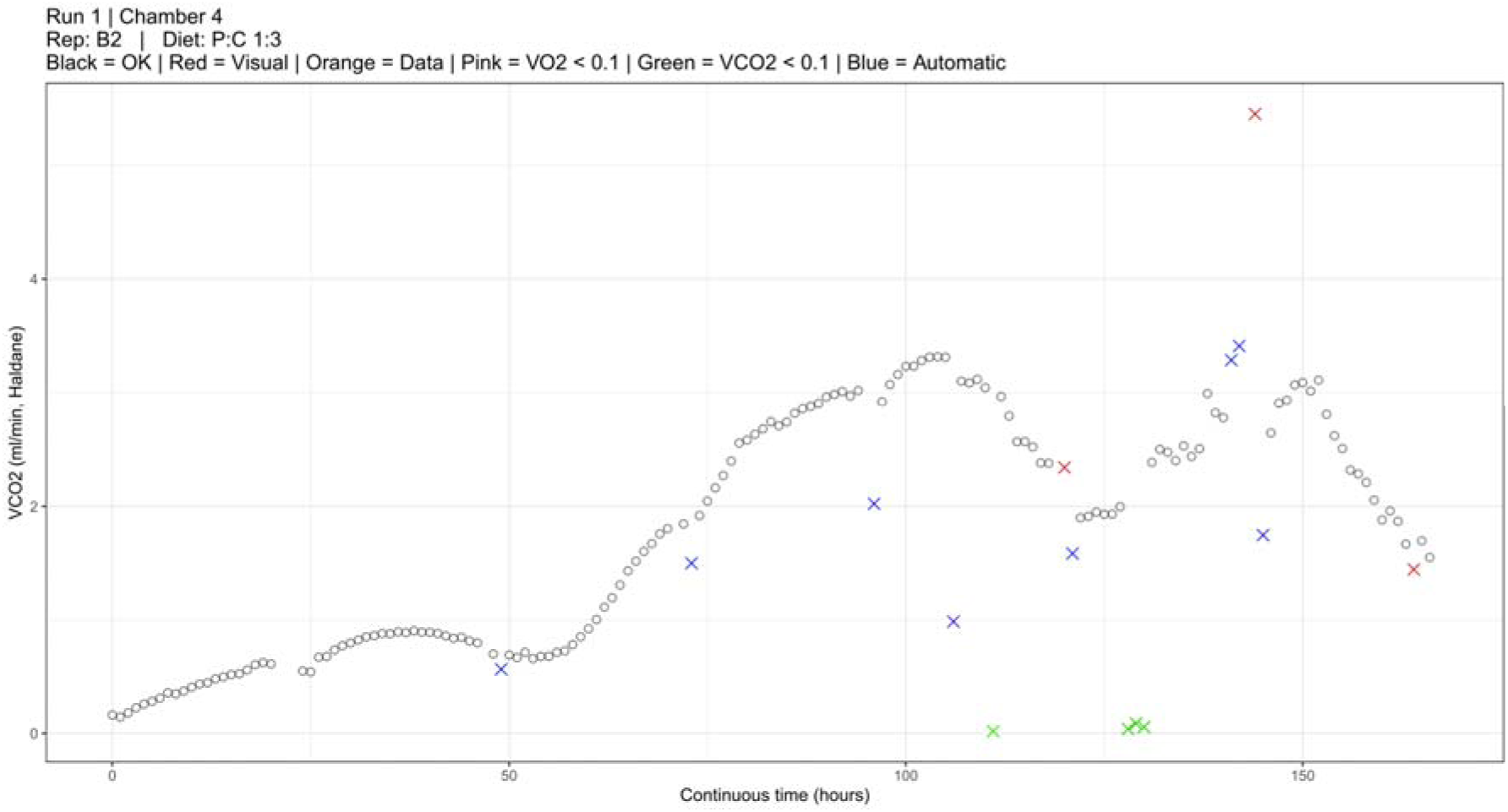
Rate of CO_2_ production (mL min^-1^) for chamber 4 (replicate B2, P:C 1:3 diet) during experimental week 1, illustrating measurements excluded by visual screening (red), low-signal criteria (green), and automatic outlier detection (blue). Accepted measurements are shown in black.

## Notes

### Competing Interest Statement

The authors have declared no competing interest.

### Summary of Updates

We have recalculated metabolic data to adjust for volume changes when RER is different from one, which has changed values only slightly. We have also updated the text according to reviewer comments.

